# ARYANA-BS: Context-Aware Alignment of Bisulfite-Sequencing Reads

**DOI:** 10.1101/2024.01.20.576080

**Authors:** Hassan Nikaein, Ali Sharifi-Zarchi, Afsoon Afzal, Saeedeh Ezzati, Farzane Rasti, Hamidreza Chitsaz, Govindarajan Kunde-Ramamoorthy

## Abstract

**Motivation:** DNA methylation plays a crucial role in biological processes, including imprinting, development, inflammation, and several disorders, such as cancer. Bisulfite sequencing (BS) is the gold standard for single-base resolution in measuring DNA methylation. This process involves treating genomic DNA with sodium bisulfite, followed by polymerase chain reaction (PCR), converting unmethylated cytosines to thymines (C to T) and guanines to adenines (G to A). However, aligning reads obtained through next-generation sequencing (NGS) of the converted DNA is challenging due to the high number of mismatches caused by these conversions.

Various BS aligners aim to simplify BS read alignment to conventional DNA alignment by modifying the reference genome and/or reads. Methods include three-letter alignment and wild-card alignment, each with its limitations.

This work introduces a novel BS aligner, ARYANA-BS, which departs from conventional DNA aligners by considering base alterations in BS reads within its alignment engine. Leveraging well-established DNA methylation patterns in different genomic contexts, ARYANA-BS generates five indexes from the reference, aligns each read to all indexes, and selects the hit with the minimum penalty. To further enhance alignment accuracy, an optional EM step has been developed, incorporating methylation probability information in the decision-making process for the index with the minimum penalty for each read. The presented approach seeks to improve the accuracy of BS read alignment by accounting for the intricacies of DNA methylation patterns in diverse genomic contexts.

**Results:** Our experimental results, based on both simulated and real data, demonstrate that ARYANA-BS achieves state-of-the-art accuracy while maintaining competitive speed and memory usage.

**Availability:** The source code of ARYANA and ARYANA-BS, the read simulator for both normal and bisulfite-treated reads, SAM file analyzer which is used for post processing of the alignment penalties, and test procedures for benchmarking different aligners using simulated and real data, are publicly available in **https://github.com/hnikaein/aryana**.

**Supplementary information:** Supplementary data are available at *Journal Name* online.

## Introduction

DNA methylation is the earliest recognized epigenetic mechanism that is found primarily in vertebrates. This process involves the attachment of a methyl group to carbon 5 of genomic cytosine, resulting in 5-methylcytosine [36]. DNA methylation plays a crucial role in a multitude of biological processes, including but not limited to imprinting, differentiation, development, inflammation, transcriptional silencing, and various diseases such as cancer [35]. Therefore, analyzing and evaluating methylation data can provide extensive information about the regulatory mechanisms of vertebrate cellular life.

Most methylated cytosines are found adjacent to a guanine nucleotide, forming a CpG site. However, cytosines in the forms of CpA, CpT, and CpC are typically not methylated [36]. Many regions of a vertebrate genome lack CpGs due to the deamination of methylated cytosines, which convert them to uracil. During DNA replication, uracil is replaced by thymine, resulting in a C to T conversion [14]. Specific regions in the vertebrate genome, called CpG islands, have a higher ratio of CpGs and GC content than other genomic regions. CpG islands are usually hypomethylated (i.e., having lower DNA methylation levels), while isolated CpGs located outside of CpG islands tend to be hypermethylated (i.e., having higher DNA methylation levels) [36].

Several experimental techniques are used to measure DNA methylation levels across the genome, including methyl-DNA immunoprecipitation (MeDIP), methylation-sensitive restriction enzymes (MSREs), and bisulfite sequencing (BS) [20]. Among these techniques, BS is widely used due to its ability to provide single-base resolution and high-throughput scale when used with next-generation sequencing (NGS) technology. Treating genomic DNA with sodium bisulfite results in the deamination of unmethylated cytosines, which are converted to uracils. During polymerase chain reaction (PCR), uracils are replaced by thymines, resulting in a C to T conversion. Meanwhile, methylated cytosines remain unchanged. Using NGS, we can align reads to the reference genome and identify C to C versus C to T conversions to count the number of methylated versus unmethylated cytosines, respectively.

It is important to note that the reverse complement strand obtained by PCR shows G to G versus G to A conversions in positions opposite to methylated versus unmethylated cytosines, respectively (Figure 1).

**Fig. 1.**
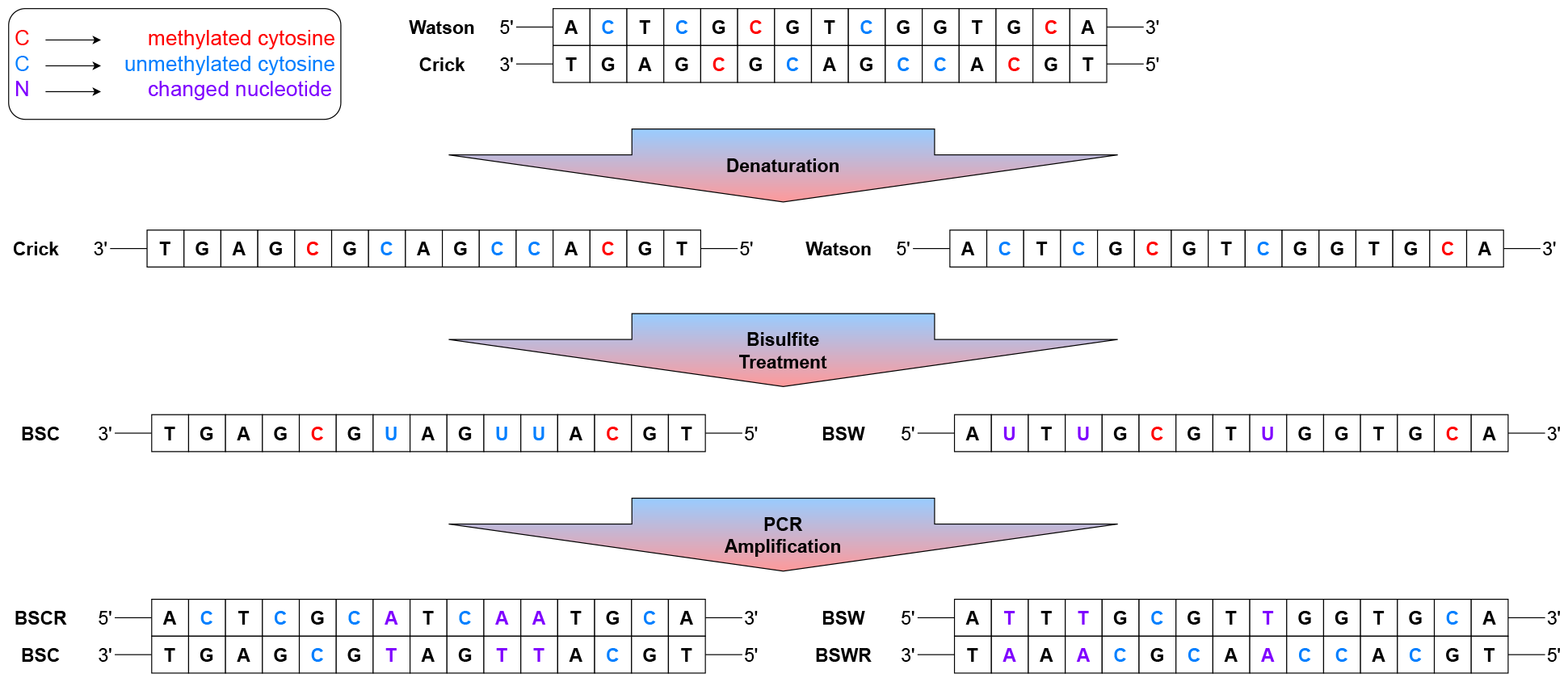
The bisulfite sequencing procedure. In the first step, an enzyme unwinds the DNA into Watson and Crick fragments. Then, during bisulfite treatment, the unmethylated cytosines are converted to uracil. Furthermore, during PCR, all uracils are converted to thymines in the final step. This conversion converts the related Crick(Watson) nucleotide from guanine to adenine.

Depending on the biological inquiry at hand, there are several sequencing methods available for profiling cytosine methylation patterns in DNA, including whole genome bisulfite sequencing (WGBS), reduced representation bisulfite sequencing (RRBS), and targeted bisulfite sequencing. WGBS involves the treatment of the entire genomic DNA with sodium bisulfite, followed by PCR amplification and sequencing, resulting in a comprehensive view of cytosine methylation patterns across the entire genome, albeit at a higher NGS cost. In contrast, RRBS targets CpG-rich regions such as promoters and CpG islands by using specific methylation-insensitive restriction enzymes, enabling the generation of more data at a lower NGS cost [23]. Lastly, targeted bisulfite sequencing is employed when we want to analyze the methylation status of a specific gene or genomic region.

A major challenge associated with analyzing WGBS and RRBS data involves the alignment of NGS reads to the reference genome. In conventional DNA short read aligners such as BWA [22], Bowtie2 [21], and others, all C to T and G to A conversions are interpreted as mismatches. Many of these aligners rely on a seed-and-extend strategy [31], which may not be effective in finding exact matches (seed sequences) when mismatches occur frequently. Therefore, specific aligners are needed for the analysis of bisulfite sequencing reads.

Many BS short read aligners use the primary strategy of adapting BS reads and/or the reference genome to enable conventional short read aligners to align them without modification. However, two different methods have emerged to resolve the C to T conversions (G to A conversions can be resolved in a similar manner).

The first method is the three-letter alignment, which involves converting all Cs of both reference and reads to Ts, thereby eliminating mismatches due to bisulfite C to T conversions and making it easy to use a conventional short-read aligner [10]. Bismark [18] produces two indices from the reference genome by converting Cs to Ts and Gs to As and then aligns the reads to each of them using the same conversions. This adaptation of the data, however, results in the loss of the distinctions between Cs and Ts in both reference and reads, and consequently, read Cs can be aligned to genomic Ts, which is not possible in bisulfite conversion. This information loss leads to multiple alignment hits across the reference at the same score, reducing the number of reads with a unique alignment position [4]. Some methods have extended this approach to introduce a two-letter alignment by simultaneously converting Cs to Ts and Gs to As in both reference and reads [12], but this approach leads to even more information loss than the three-letter method.

The second method is wild-card alignment, which involves replacing all Cs of the reference with the wild-card letter Y, which can stand for either C or T in IUPAC. In this way, Cs and Ts of the reads can be aligned to the reference Ys. BSMAP [38] uses a hash table to record all k-mers in the reference genome and records all C-to-T variations of the reference k-mers to implement the wild-card alignment. Unlike the three-letter alignment, the wild-card alignment does not result in information loss. However, it has a bias toward aligning reads from hypermethylated genomic regions better, as reads coming from such regions have more Cs, which can only be mapped to reference Ys. In hypomethylated regions, Cs diminish after C to T conversion, and Ts of a read can be aligned to both Ys and Ts of the reference, leading to nonunique alignment. Many downstream analyses, including the estimation of the methylation ratio of each genomic cytosine, eliminate nonuniquely aligned reads. As a result, this method suffers from a systematic overestimation of the methylation ratio, since reads from hypermethylated regions have a better chance of unique alignment [4].

In this article, we introduce ARYANA-BS, a novel short read aligner for bisulfite sequencing that builds upon our previous work on the conventional short read aligner, ARYANA [11]. Unlike many existing methods that rely on adapting reads and/or references, ARYANA-BS incorporates specific treatments for BS reads during the alignment process. Specifically, we have enhanced the methods for gap-filling, computing edit distance, and alignment score (penalty), considering the possible base conversions that may occur due to bisulfite treatment, based on the distinct contexts in which they occur. These contexts include differentiating between CpG and non-CpG cytosines, as well as CpG island and non-CpG island regions. Additionally, we utilize five distinct indexes for optimizing the seed-finding procedure, as described in detail in the Methods section. Furthermore, we developed an optional EM step that leverages the methylation probability of each cytosine to improve the final alignment result.

We benchmarked ARYANA-BS, as well as several widely-used BS aligners, including Bismark [18], BSMAP [38], bwa-meth [27] (a three-letter aligner), BSBolt [9] (another three-letter aligner), and abismal [7] (a two-letter aligner), using both simulated and real data. We utilized a pair of simulators, one developed by us and another introduced in independent research, and found consistent results. Additionally, we used real data that had previously been used in independent benchmarks of BS aligners. We provide time, memory, and accuracy comparisons of the aligners, and show that ARYANA-BS outperforms the other aligners in terms of alignment accuracy on simulated and real data, while still maintaining competitive speed and memory usage.

## Results

We conducted a comparative analysis of ARYANA-BS against five other widely used WGBS aligners: BSMAP, Bismark, bwa-meth, BSBolt, and abismal. BSMAP was chosen for its acceptable results in prior studies, while Bismark is known for its effectiveness and widespread use. bwa-meth is an adaptation of BWA mem, while BSBolt is a new aligner, and abismal employs a unique approach of two-letter mapping. The default configuration parameters were employed for all tools. The comparison was carried out using SAM file analyses, with hg38 as the reference and the CpG islands file obtained from http://hgdownload.cse.ucsc.edu/goldenpath/hg38/database/cpgIslandExt.txt.gz.

In the initial phase, we generated simulated bisulfite-treated reads from the hg38 genome using both the ARYANA and SHERMAN read simulators [1]. In the subsequent phase, we conducted an extensive evaluation of our algorithm using real data sourced from SRR3469520, which is available from the NCBI. This real-world dataset provided a valuable benchmark for assessing the efficacy of our methodology. Finally, in the last phase of our investigation, we delved into the impact of incorporating an additional expectation-maximization (EM) step on the outcomes generated by ARYANA-BS. This step was introduced to further refine the results and explore potential enhancements to the algorithm’s efficacy.

It should be noted that although Bismark supports paired-end alignment, we encountered several unsuccessful attempts to run it on paired-end data, resulting in no output. Furthermore, BSMAP can only align reads that are 144 bp or shorter, rendering it unusable in some analyses.

All tools were employed in single-threaded mode, with an AMD Ryzen 5 3600X CPU and a maximum of 48 GB memory.

### simulated data

In the initial stage, our objective is to compare ARYANA-BS with other tools based on simulated data. The data generated for this purpose were simulated using both the ARYANA simulator and Sherman. We used the ARYANA simulator to generate both 100 bp and 300 bp single-end and paired-end reads. For Sherman, we only generated 100 bp single-end reads. However, despite running for 163 hours, Sherman failed to generate any output for the simulation of 300 bp single-end reads.

The ARYANA simulator was developed to generate reads for regular sequencing as well as bisulfite sequencing. To simulate bisulfite sequencing, the simulator takes the reference genome as input and produces SNPs after loading the reference genome into memory. During the read generation process, the simulator takes into account the insertion, deletion, and mismatch ratio provided in the input. Moreover, the simulator generates reads from both the positive and negative strands. In the case of bisulfite sequencing, the simulator also incorporates the effects of negative strand and PCR in the output reads. These effects are based on G-to-A and C-to-T conversions that are dependent on the C or G in the reference genome. The CpG islands from the input are utilized by the simulator to make bisulfite conversions based on the context of cytosines. The default methylation ratios for the CpGs inside the CpG islands are set at 0.1, while outside the CpG islands the default is 0.9, and for the CpH, the default methylation ratio is 0.01.

The primary aim of the first comparison is to evaluate the accuracy and performance of various tools. In the subsequent two comparisons, we closely examine the effect of possible biases in the genomic context on the performance of the tools.

### Comparison of Memory, Time, False Discovery Rate (FDR), and Correctly Mapped Reads Percentage

In this analysis, we evaluated the accuracy of the tools based on simulated reads. We generated 1,000,000 reads for each simulation and used the tools to align them with the reference genome. We used the ARYANA simulator for four simulations that differed in the length of reads and whether they were single-end or paired-end sequencing. The ARYANA simulator was configured to use 4 million single-nucleotide polymorphisms (SNPs). Additionally, the Sherman simulator was used to simulate 100 base pair (bp) single-end reads.

We considered an alignment to be correct if the reported position by the tool for the alignment differed by no more than one base pair from the actual position reported by the simulator. We also computed the FDR using the following formula:

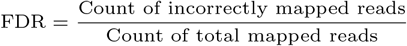

In all simulation data generated by ARYANA and the Sherman simulator, ARYANA-BS outperformed all other tools in terms of the number of correctly mapped reads. However, BSMAP had comparable results in three simulations, but it only supports reads shorter than 145 bp, making it unsuitable for sequencers of larger read sizes.

The best whole-genome bisulfite sequencing (WGBS) aligner in terms of performance analyses was abismal, with the minimum runtime and memory usage of all tools. However, its accuracy was not acceptable. Bismark had better memory usage, but its accuracy was worse than that of any other tool. ARYANA-BS had comparable runtime to other tools, and in simulations with longer read sizes, it was faster than others. Moreover, its memory usage stood at an average level, closely aligning with that of BSMAP.

Although the FDR of ARYANA-BS was not as low as that of bwa-meth or BSBolt, we should work on reducing it. (see Figure 2)

**Fig. 2.**
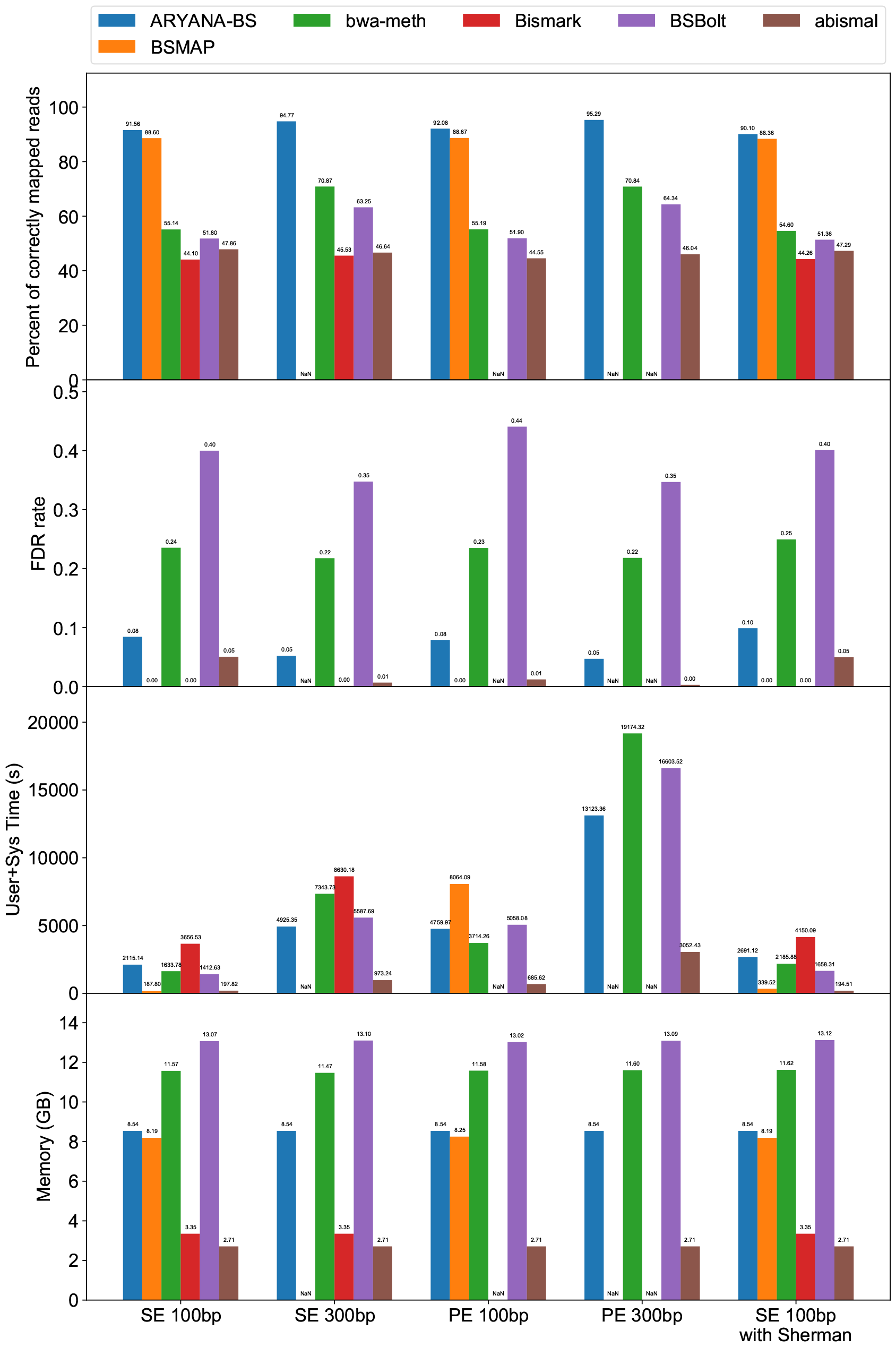
Comparison of ARYANA, BSMAP, bwa-meth, Bismark, BSBolt, abismal. We have shown four comparisons: percent of correctly mapped reads, FDR (false discovery rate), memory footprint, and runtime. We compared these tools with five simulated inputs: 100 base pairs single-end reads, 300 base pairs single-end reads, 100 base pairs paired-end reads, 300 base pairs paired-end reads, all simulated with ARYANA read simulator, and 100 base pairs single-end reads simulated with Sherman.

In Figure 3, we demonstrated that most of the reads aligned to the context-aware indexes instead of the original index and three-letter indexes. This result highlights the importance of context-aware alignment in the improvements of ARYANA-BS.

**Fig. 3.**
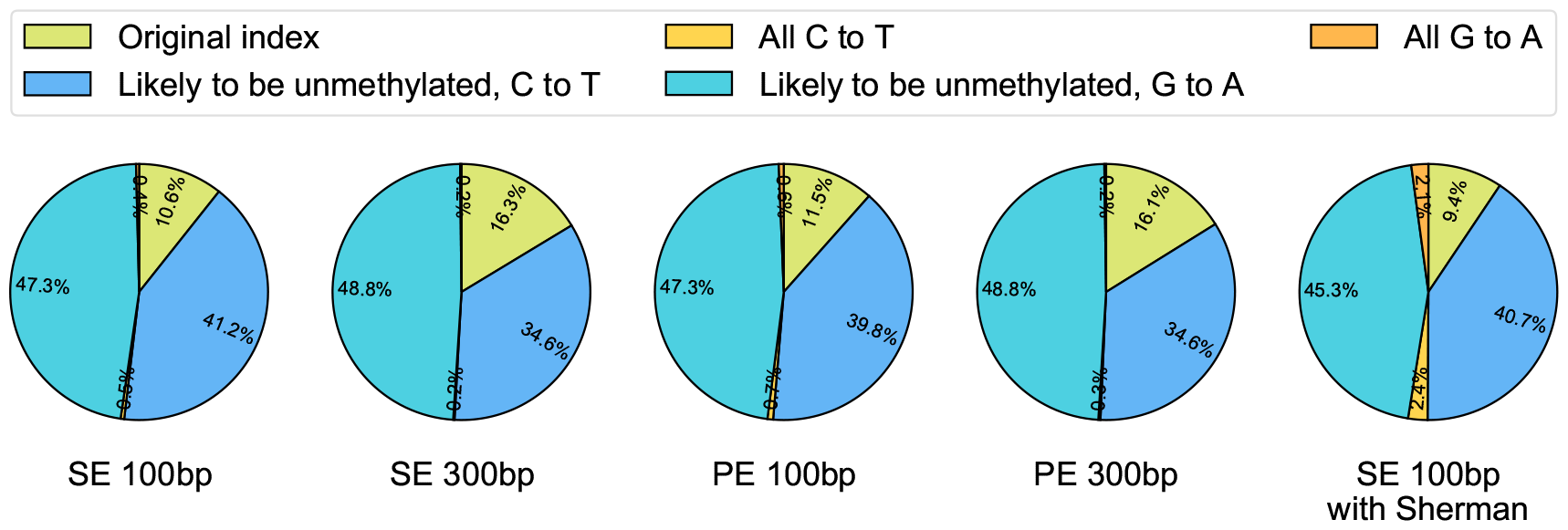
Percent of alignment of reads to each index. This figure shows the percentage of each dataset that aligned to each index. The majority of the reads aligned to the context-aware indexes, rather than the original index and three-letter indexes.

### Single Nucleotide Polymorphisms (SNPs) and Indel Rates

In Figure 4, we present a comparison of correctly mapped reads for simulated reads with varying SNP counts and different indel rates. This analysis showcases the efficacy of the tool in genomic change analyses, such as cancer analyses.

**Fig. 4.**
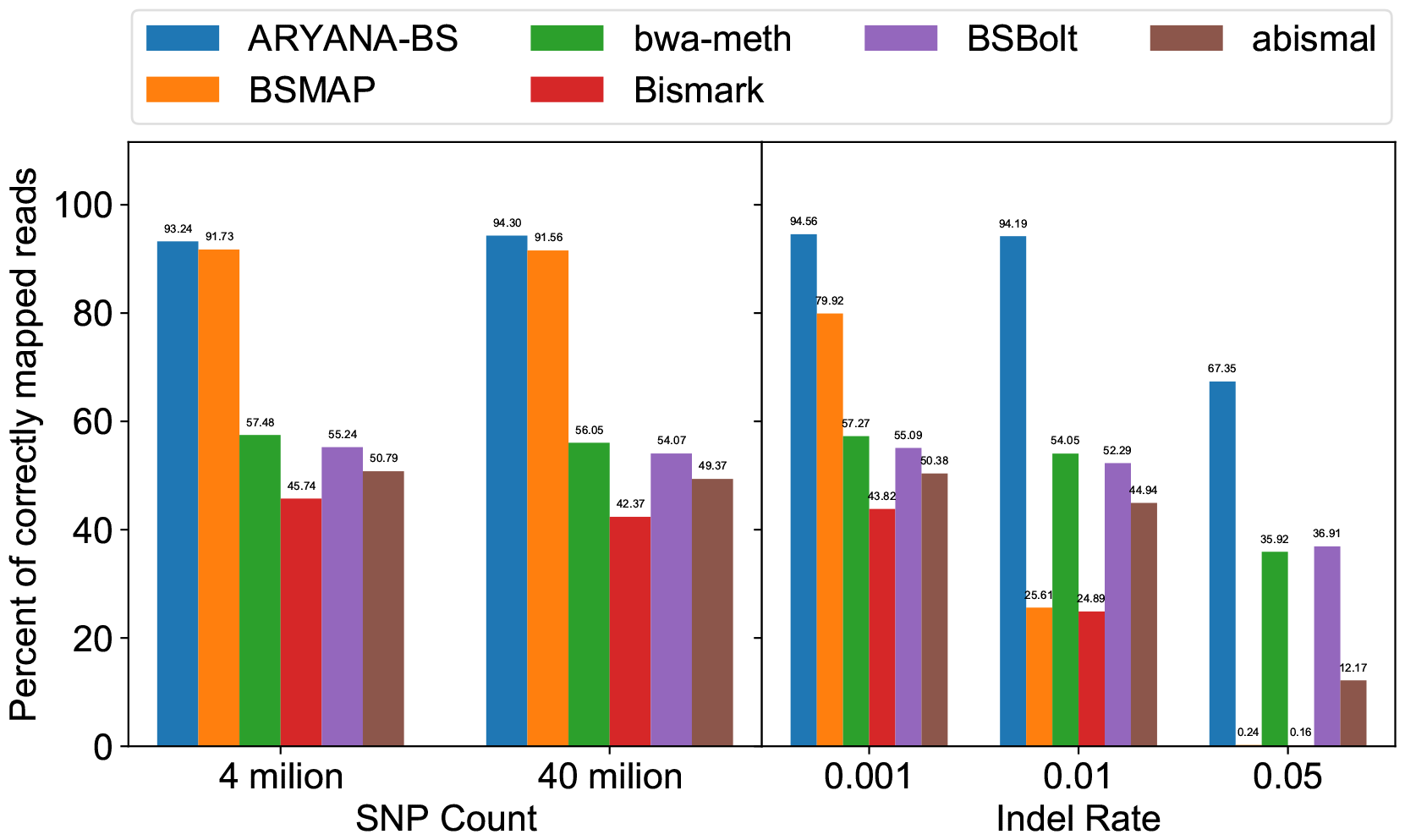
Scrutinize of tools against different SNP counts and indel rates. We generated two different simulated reads with different SNP counts. This figure shows the percentage of correctly mapped reads for each tool case. In both cases, ARYANA-BS performed better than the other tools. Additionally, we generated three different simulated reads with different indel rates, and again ARYANA-BS mapped more reads correctly with different indel rates than others.

In the initial simulation, we simulated two sets of reads with SNP counts of 4 million and 40 million. Although this change has a negligible impact on the tool’s performance, it is worth noting that ARYANA-BS outperforms all other tools in both tests.

To further investigate ARYANA-BS’s performance concerning the sequence with nucleotide insertion or deletion, we simulated three sets of reads with varying indel rates. Specifically, we simulated reads with a low indel rate of 0.001, an indel rate of 0.01, and a high indel rate of 0.05. As illustrated in Figure 4, even a minuscule indel rate of 0.001 significantly impacts BSMAP’s performance. Additionally, the results from 0.01 and 0.05 indel rates indicate that ARYANA-BS is more resistant to indel rates than other tools, and it consistently aligns more reads correctly than any other tool.

### Percentage of Reads in CpG Islands and Methylation Percentage

We conducted three simulations to investigate the bias of ARYANA-BS toward reads from CpG islands. In the first simulation, no reads originated from CpG islands. In the second simulation, 50% of the reads came from CpG islands, while all the reads in the third simulation belonged to CpG islands. Although the results were similar across all simulations, ARYANA-BS consistently outperformed the other tools (see Figure 5).

**Fig. 5.**
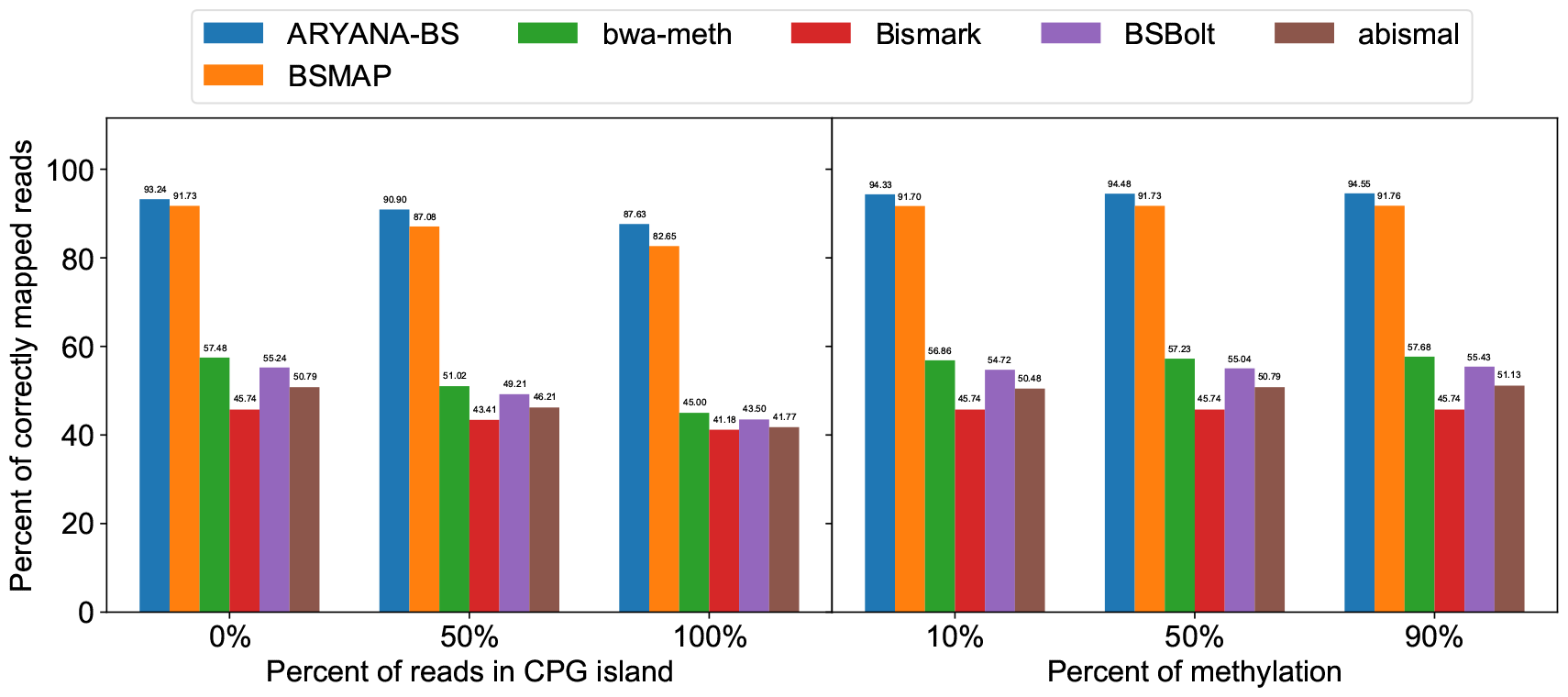
Scrutinize of tools against different percent of reads in CpG island and methylation. This figure shows the performance of each tool for different percentages of reads in CpG islands and the percent of methylation in the simulated reads created with the ARYANA read simulator. ARYANA-BS is better than other tools in all cases.

We further analyzed the bias of ARYANA-BS and other tools toward methylation percentages of reads. To accomplish this, we simulated reads with varying degrees of methylation, including hypomethylated reads (10% methylation), averagely methylated reads (50% methylation), and hypermethylated reads (90% methylation). In all three cases, ARYANA-BS exhibited superior performance compared to other tools. However, no significant differences were observed among all the tools (see Figure 5).

### Real Data Analysis

To evaluate the accuracy of the proposed algorithm on real data, we conducted a comparison of the alignment efficiency between ARYANA-BS and other available tools. The first one million reads from each file of SRR3469520 run from NCBI (https://trace.ncbi.nlm.nih.gov/Traces/index.html?view=run_browser&acc=SRR3469520) were used for this purpose.

As we lacked knowledge of the exact position of each read in the real data, we computed the alignment penalty of each read for each tool to obtain a fair comparison between the two tools. To ensure a level playing field, we utilized penalty scores from bwa-meth. Mismatch penalties were set at 4, gap open penalties were set at 6, gap extension penalties were set at 1, and match scores were set at 4. The difference between the alignment penalty of the two tools was then computed for each read.

Figure 6 showcases the count of reads with differentiations in comparison between ARYANA-BS and other tools, as well as the number of reads each tool could align while the other failed to do so. It is noteworthy that we did not employ Bismark because it was unable to generate output in the case of paired-end reads. It is evident that ARYANA-BS outperformed other tools in this analysis. Although BSMAP demonstrated lower alignment penalty for 4% of reads, it failed to align approximately 29.4% of reads.

**Fig. 6.**
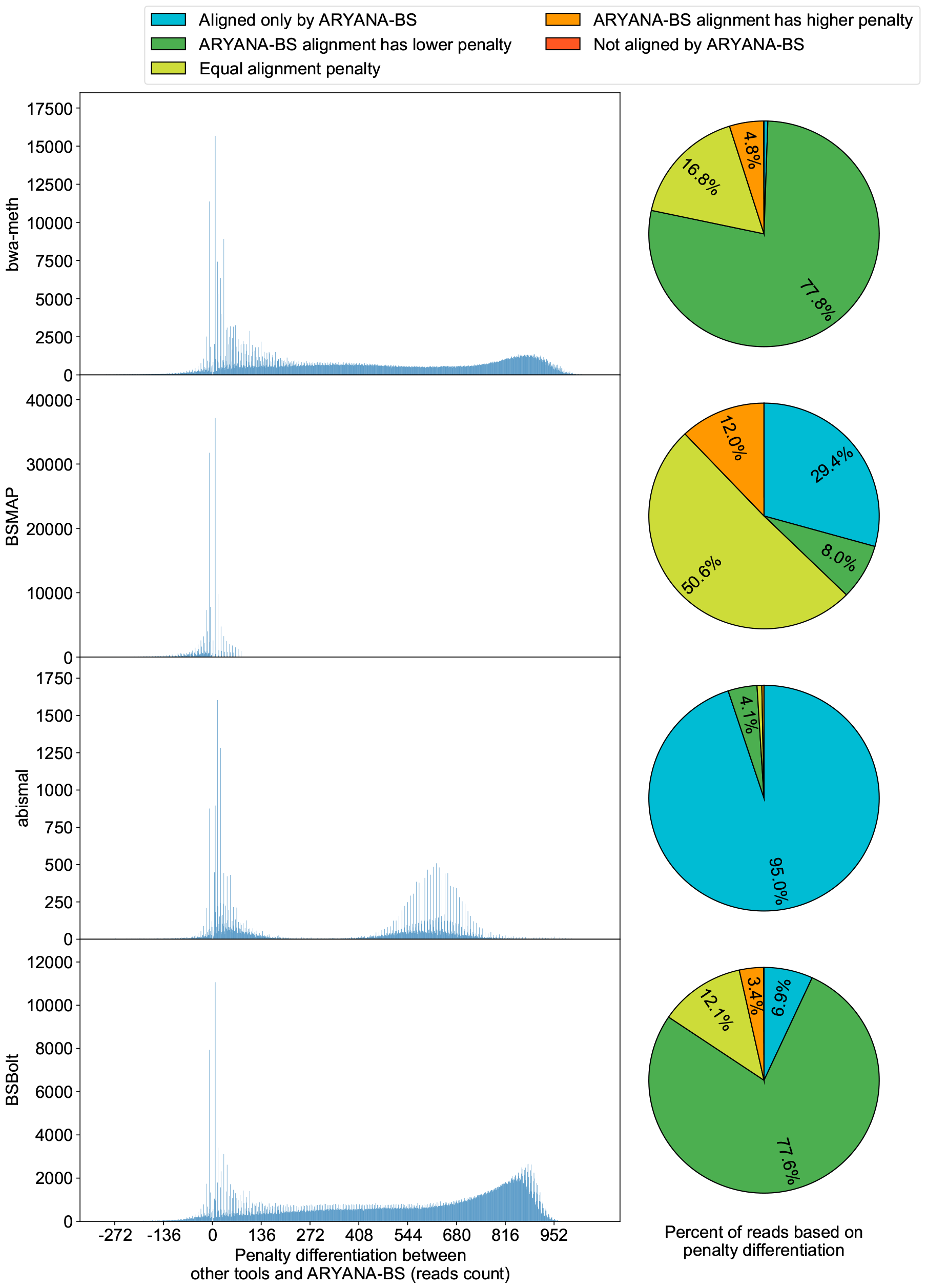
Comparison of ARYANA-BS against bwa-meth, BSMAP, abismal, and BSBolt with real data. We used the read alignment penalty to compare the tools with real data to determine the alignment quality of each tool. For this comparison, we have calculated the penalty for each tool on each read and the differentiation between the ARYANA-BS alignment penalty and the other tool’s penalty. In this figure, we have indicated the count of reads with each differentiation and the number of reads that were aligned only by ARYANA-BS or the other tool. Note: the box chart was trimmed to avoid large plots. Additionally, the reads with equal alignment penalties omitted from the box chart.

### Analysis of the additional EM-step

To investigate the impact of the EM (Expectation-Maximization) step on the improvement of ARYANA-BS results, a dataset consisting of 5 million reads, each with a length of 100 base pairs, was generated from chromosome 21. This was done to achieve an approximate sequencing depth of approximately 10X, considering resource constraints that prevented a genome-wide analysis.

A summary of the ARYANA-BS execution, both without the EM step and with the EM step applied to this dataset, is presented in Table 1. In summary, the implementation of the EM step was able to reduce the ARYANA-BS error by 3.23% without any change in the needed memory, incurring an additional 9.76% runtime. Furthermore, it increased the percentage of reads aligned to their correct positions from 90.73% to 91.03%.

**Table 1.** Analysis of the additional EM-step on ARYANA-BS.

|  | Percent of correctly mapped reads | User+Sys Time (s) | Memory (GB) |
| --- | --- | --- | --- |
| ARYANA-BS | 90.73 | 3634.33 | 1.87 |
| ARYANA-BS with EM-step | 91.03 | 3991.57 | 1.87 |

## Methods

As mentioned in the background section, the current paradigm in many BS short read aligners is to use a conventional short read aligner of untreated genomic DNA as a black box (unchanged or with minimal changes) and, rather, adapt the reference genome and/or reads (i.e., wild-card, two-letter or three-letter). Information loss, as observed in two- or three-letter methods, as well as a biased alignment rate depending on the methylation ratio of the genomic region in wild-card aligners, has been extensively discussed in the literature as the major limitation of these methods [19].

### Context-aware paradigm

In ARYANA-BS we tried to overcome shortcomings of the prior methods by developing a new paradigm that we call *context-aware alignment*. In this paradigm, we try to adapt the seed-and-extend algorithm to better match the BS short read alignment. For this purpose, we need to take well-known biological information about the patterns of DNA methylation in different contexts into account:

- *Dinucleotide context:* The distribution of the DNA methylation ratio of cytosines present in CpG dinucleotides is significantly different from the other contexts (i.e., CpA, CpC and CpT dinucleotides). Although the presence of 5-methylcytosines in DNA was discovered in the 1940s, there was very limited knowledge about non-CpG methylation by the early 2010s [25]. Following major developments in DNA-methylation assays, particularly RRBS and WGBS, more studies have shown the existence of non-CpG methylation in some human tissues (e.g., brain), as well as embryonic stem cells [34]. Further research showed biological functions of non-CpG methylation [26, 17, 28], and suggested epigenetic mechanisms that regulate the presence of non-CpG methylation [2, 3]. Some packages perform downstream analysis of DNA methylation data (e.g., finding differentially methylated regions) by distinguishing between CpG vs. non-CpG methylation [5]. However, the extreme alteration of methylation distribution in CpG vs. non-CpG context is considered neither in two/three letters, nor in wild-card alignment.
- *Genomic context:* A significant alteration of the methylation pattern was observed in different genomic contexts, particularly in active regions (e.g., promoters of expressed genes) [8, 13] vs. silenced regions (e.g., heterochromatin) [30, 33, 29]. These alterations have been confirmed at single-cell, cell-type resolution [40]. A common observation has been a significant alteration of CpG methylation distribution in CpG islands: although isolated CpGs tend to be hypermethylated in many tissues, those present in CpG-rich regions such as CpG islands, particularly in proximity to active promoters, show lower methylation levels [15, 32].

We should note that these rules are not zero-one rules, as there are well-known hypermethylated CpG islands in different tissues and diseases [16, 39, 37, 6]. There are also observations about flanking regions of CpG islands, called CpG island shores [24]. Altered methylation distribution of isolated CpGs vs. those present in CpG islands can be employed as important prior information that was not taken into account in two/three letter or wild-card alignment.

ARYANA-BS is an enhancement of our conventional short-read aligner ARYANA [11], which performs context-aware alignment of BS reads. To better describe ARYANA-BS, we need to have a quick glance over how ARYANA works, and then review what enhancements are made to build ARYANA-BS.

### A review of ARYANA

ARYANA uses a seed-and-extend paradigm for aligning short reads of genomic DNA. It creates a Burrows-Wheeler Transform (BWT) index of the genome using the BWA engine [22], partitions the reference genome into equal-sized windows, and finds maximal substrings that exactly match between the read and the reference (seeds). For each seed, it scores the genomic windows that contain the seed. It then selects a subset of windows with the maximum scores, finds the precise alignment penalty of the read in each window, and reports the alignment position with the minimum penalty. We introduced a bidirectional BWT search algorithm and used it for seed-extension in both directions. The algorithm creates a BWT index on the reference and its reverse complement, as well as another BWT index on the reverse of the reference and the reference complement. We used the former index to find a minimal seed that did not overlap with the previous seed and used the latter index to extend it. This allowed us to make a significant speedup by creating non overlapping seeds over the read instead of searching every base of the read for a new seed. Furthermore, several algorithmic enhancements and techniques are used to improve the efficiency of ARYANA. For example, it uses a reset-free hash table as the data structure to search and update window scores. By “reset-free,” we mean that the hash table does not need to be cleared for each new read. ARYANA is multithreaded and can process both single-end and paired-end reads.

ARYANA-BS employs context awareness of DNA methylation patterns in both seed finding and extension phases of read alignment, as described below:

### Context-aware seed finding

By considering alterations in the cytosine-methylation probability distribution across dinucleotide and genomic contexts, ARYANA-BS creates five different indexes from the reference genome (figure 7):

- *Index a:* the original reference genome, unchanged.
- *Index b:* all cytosines that are more likely to be unmethylated, according to their dinucleotide and genomic contexts, are converted to thymines. This resembles the most probable sequence of the genome, after being treated with sodium bisulfite. The default strategy is to consider CpGs outside CpG islands more likely to be methylated, and all other cytosines (e.g., CpAs, CpCs, and CpTs across the genome as well as CpGs inside CpG islands) more likely to be unmethylated. The list of genomic regions considered to be hypomethylated is provided as an input to the algorithm. By default, it contains CpG islands, but it can be changed depending on the specific organism, tissue or cell type to contain most likely hypomethylated regions across the whole genome.
- *Index c:* all cytosines are converted to thymines, regardless of their genomic or dinucleotide contexts. This index is identical to the one used in the three-letter strategy, but as we later show, the extension phase of ARYANA-BS is different from what is used in three-letter algorithms.
- *Indexes b’ and c’:* similar to genomes b and c, in which, guanine to adenine conversions are made. It is very important to note that PCR amplification of bisulfite-treated DNA fragments can lead to four different sequences from the same region, depending on whether the original forward vs. reverse fragment is used as the PCR template, and whether the final fragment being sequenced is similar or reverse complement to the template 1. The PCR product contains fragments that are reverse complemented to the original template from the genomic bisulfite-treated DNA. These fragments contain G to A conversions, which distinguish them from reverse-strand fragments of the original bisulfite-treated DNA. In genome b’, we select all Gs in the forward strand of the reference genome where the opposite C in the reverse strand is more likely to be unmethylated (as described in genome b) and convert them to A. In genome c’, we convert all Gs in the forward strand of the reference genome to A.

It is important to note that for each of the five above genomes, we concatenate the sequence and its reverse complement (with a separator in the middle) and use the resulting sequence to create the BWT index. In this way, both forward and reverse reads can be aligned.

**Fig. 7.**
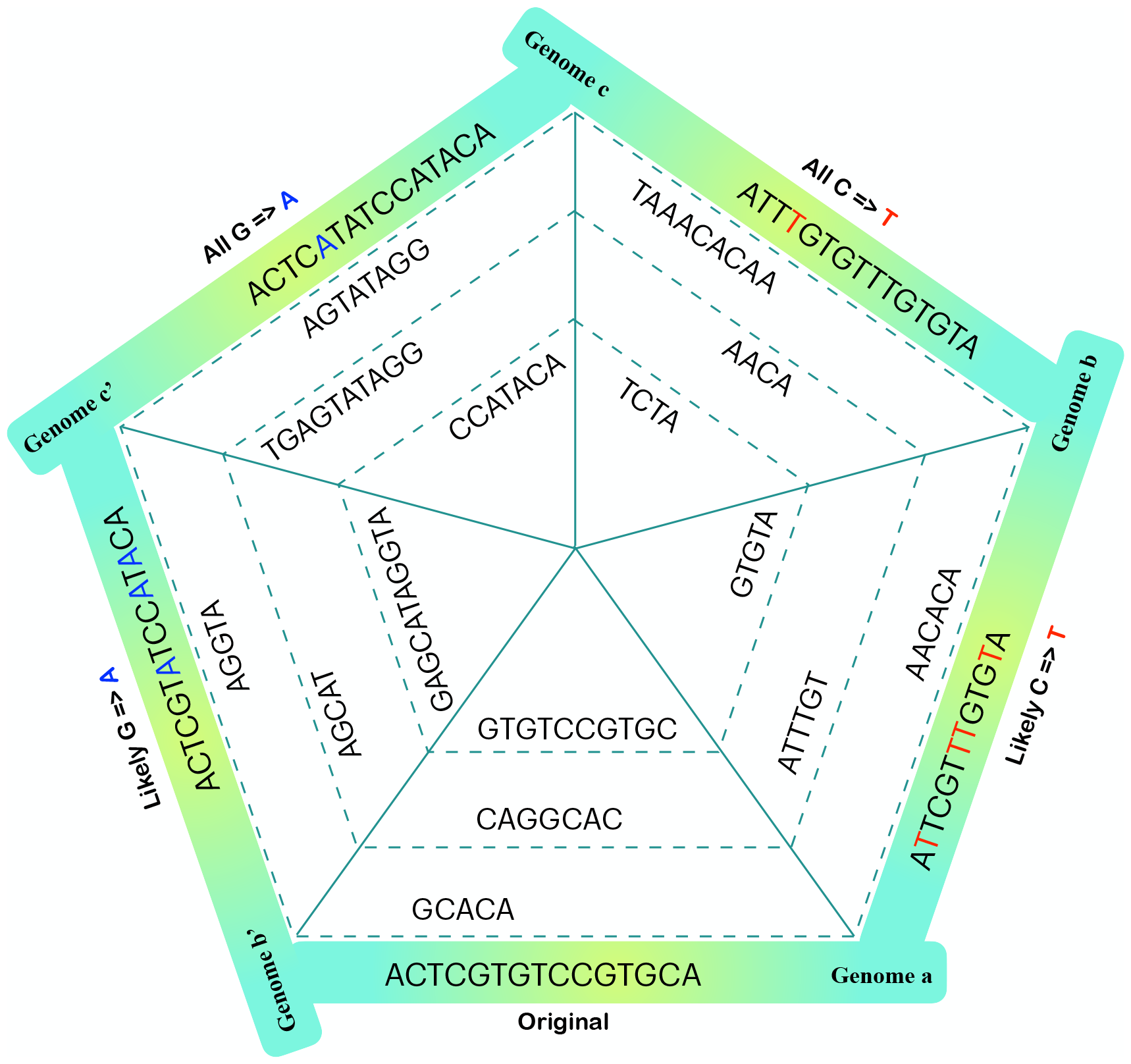
Context-aware different indexes for the ARYANA-BS seed finding algorithm. Genome a: original reference genome. Genome b: all cytosines that are expected to be unmethylated, according to the prior context information, are converted to thymines. Genome c: all cytosines, regardless of their contexts, are converted to thymines. Genomes b’ and c’: similar to genomes b and c, by converting guanines to adenines

By using these five different genomes, we ensure that the context information is used to create longer exact seeds without information loss. Reads are NOT altered neither in seed finding, nor extension.

### Context-aware seed extension

As we mentioned earlier, both ARYANA and ARYANA-BS partition each reference genome into equal-sized windows to find the best part of the genome for read mapping. Each read was aligned independently to each of the five references. For each reference, we find maximal exact matches and score the genomic windows that contain the seed (for more details, see ARYANA article [11]). Then, we list the top scored windows and try to extend individual seeds in each window to cover match the whole read.

This extension phase is mainly performed by the following steps:

- A maximal list of consequenced seeds in each window can be determined using a greedy algorithm that guarantees compatibility between the order of the read and the reference. The seeds in the read are arranged in a nonoverlapping fashion, making the greedy algorithm a viable approach for identifying a maximal list of consequenced seeds that are not overlapped in the reference and maintain the same order.
- Filling the gaps between each pair of consecutive seeds in the list. For this purpose, we have developed a dynamic programming algorithm for finding the optimal-matching between the part of the read between two seeds, and part of the reference located between the same pair of seeds. Although this algorithm has the same logic as Smith-Waterman, it has several differences. The scoring scheme is compliant with bisulfite-sequencing. Therefore, in genomes a, b, and c, we consider zero penalty for genomic cytosines that are mapped to read thymines. However, it is important to mention that genomic thymines mapped to read cytosines are treated as mismatches, which makes the ARYANA-BS extension different from either two/three-letter or wild-card aligners. Furthermore, we take the dinucleotide context of cytosine into account and treat CpG dinucleotides differently from CpAs, CpCs and CpTs in terms of the mapping penalty. For genomes b’ and c’, we consider a similar penalty scheme for genomic guanines that are mapped to read adenines.
- Finally, we sum the total match scores and mismatch/indel penalties for all gaps inside the consecutive seeds. If the data is paired end, we look for the best pair of matches, one of read-1 and the other for read-2, which are concordant according to the distancing arguments provided to ARYANA-BS. If no concordant alignment is identified, we can report discordant alignment of each read if specified in the arguments.

### EM Algorithm for Improving Methylation Probability Estimation and Read Alignment

Following the initial alignment phase, a subsequent post-processing step is undertaken for the aligned reads against each of the five reference genomes. This process involves the computation of context-aware alignment penalties for the reads within each of these five reference genomes. The alignment with the most favorable penalty score is selected, further enhancing the precision of the alignment. To augment the accuracy of this step, an optional EM algorithm is employed (Figure 8).

**Fig. 8.**
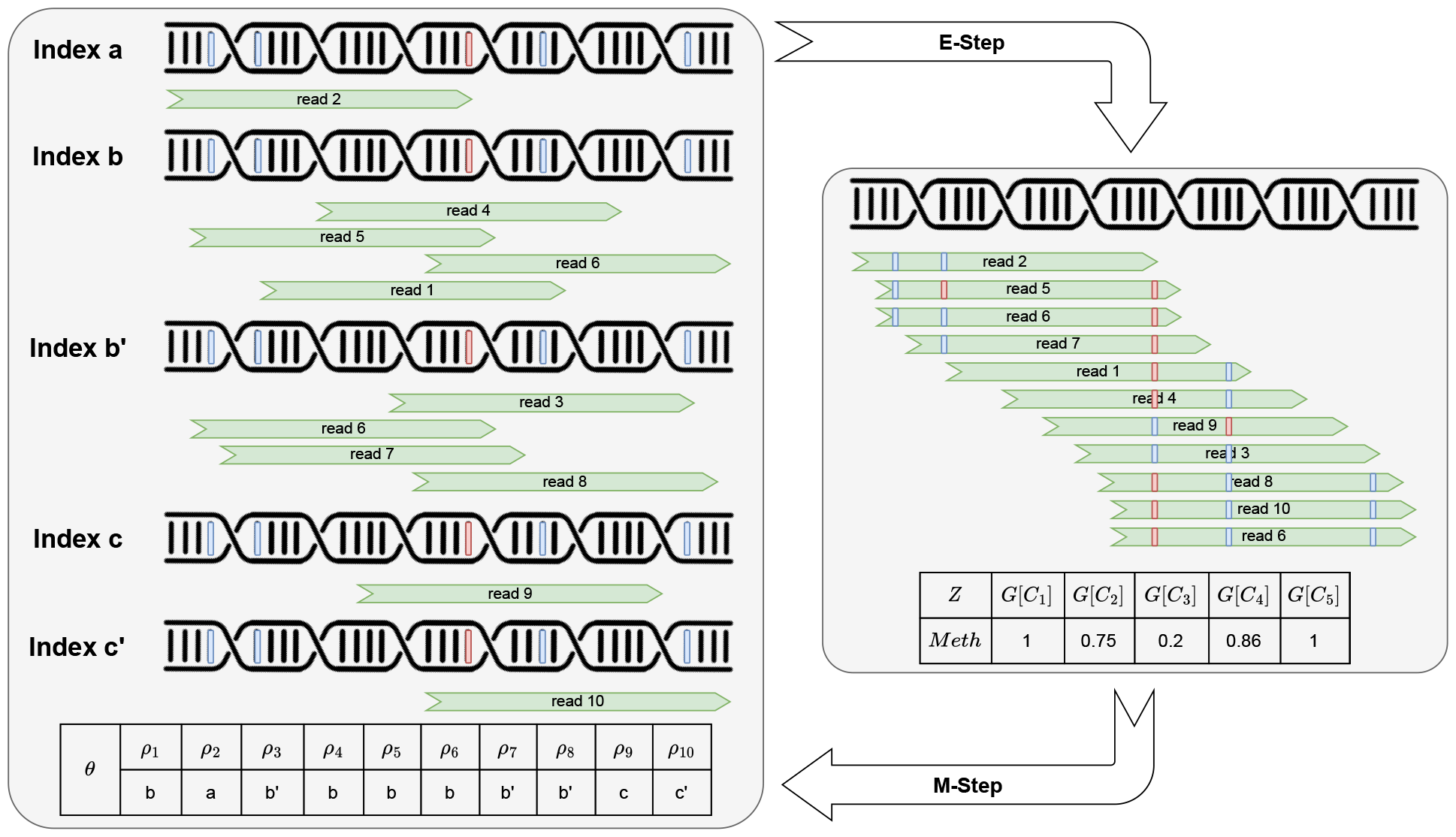
The EM-step of ARYANA-BS. In the expectation step, the probability of cytosine methylation at each site, denoted as the latent variable Z, is determined based on the alignment specified in 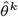 . Subsequently, in the maximization step, the best index for the placement of each read is determined based on the probability of cytosine methylation at each cytosine site, utilizing the equation 3.

The EM algorithm plays a pivotal role in estimating the methylation probability of each cytosine, treating it as a latent variable. This methylation probability is then seamlessly integrated into the alignment penalty calculation for each read within each of the five indices. This integration ensures that the resulting alignment not only optimizes the penalty score but also takes into account the methylation probability of each individual cytosine.

The algorithm introduces a variable, denoted as *θ*, which is represented as a tuple *θ* = (*ρ*_1_, *ρ*_2_, …, *ρ*_*r*_). In this representation, *ρ*_*i*_ characterizes the alignment of read *R*_*i*_ with one of the five generated indices, and *l*(*ρ*_*i*_) denotes the specific aligned location. Consequently, the dataset *X* encompasses sequences corresponding to each input read.

Furthermore, a missing variable, *Z*, is defined as follows:

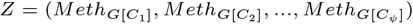

In this definition, *G*[*i*] signifies nucleotide *i* within genome *G*, and 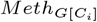 denotes the methylation state (either methylated or unmethylated) of each cytosine at the specified genomic locations.

### Expectation phase

In the E-step of the EM algorithm, we embark on the task of determining the methylation probability for each cytosine situated within CpG dinucleotides found in both the forward and reverse strands. This determination is derived from the outcomes of the read alignments against the five reference genomes. In this context, each C-to-C alignment (and G-to-G alignment for PCR reads) is considered representative of a methylated cytosine, while T-to-C and A-to-G alignments are regarded as indicative of unmethylated cytosines.

To calculate the methylation probability for each cytosine *G*[*C*_*i*_] within genome *G* (where *G*_*i*_ denotes the i-th nucleotide of genome *G*) based on the estimated parameter 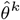, the following formula is employed:

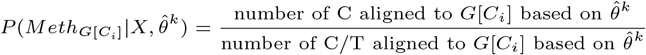

This formula calculates the likelihood of methylation for a specific cytosine *G*[*C*_*i*_] in genome *G*, taking into account the information from the read alignments and the parameter estimate 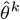 .

### Maximization phase

In the maximization step of the EM algorithm, the alignment of reads to each of the five reference indices is carried out based on the methylation probabilities computed in the expectation step. The objective function *Q* is defined as follows:

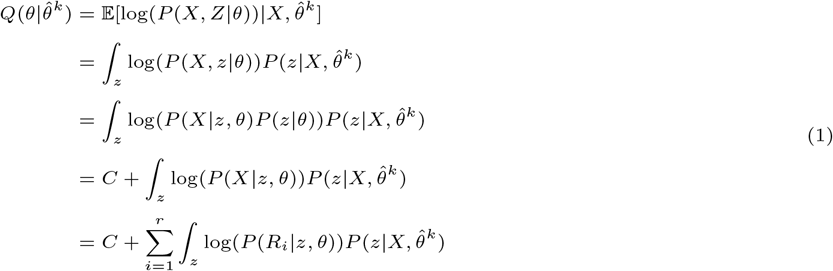

Assuming the independence of the alignment probability of *R*_*i*_[*j*] to *G*[*l*(*ρ*_*i*_) + *j*] from the alignment probability of *R*_*i*_[*j* + 1] to *G*[*l*(*ρ*_*i*_) + *j* + 1], *P* (*R*_*i*_|*z, θ*) can be computed as:

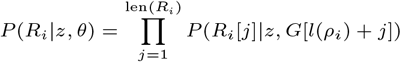

Therefore we can rewrite the expression 1 as follows:

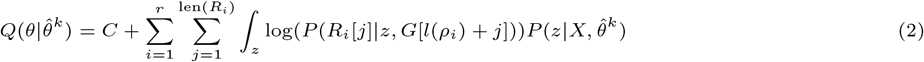

In a simplified model where *ϵ* represents the sequencer error probability, *P* (*R*_*i*_[*k*]|*z, G*[*k*^*′*^]) can be computed as:

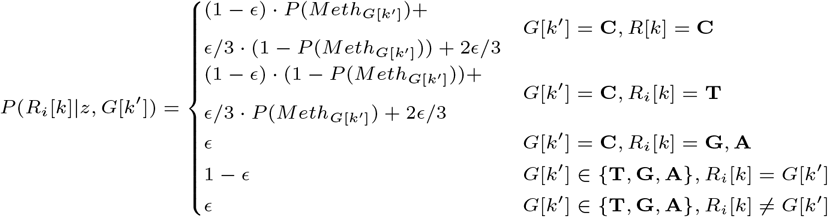

Assuming a mismatch penalty equal to *™* log(*ϵ*), log(*P* (*R*_*i*_[*k*]|*z, G*[*k*^*′*^])) can be approximated as:

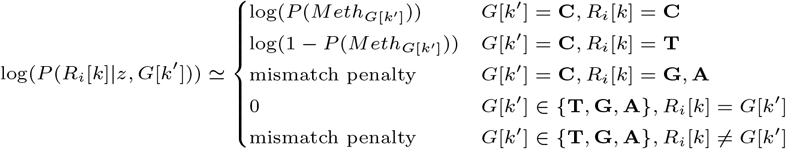

In line with the provided explanations, to determine *θ* for maximizing 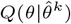, the values of *ρ*_*i*_ need to be determined in a manner that maximizes the expression 2. Given that the values of *ρ*_*i*_ are independent of each other, and the effect of each one on expression 2 is independent of the others, for each read *R*_*i*_, the value of *ρ*_*i*_ is set in a way that maximizes the following:

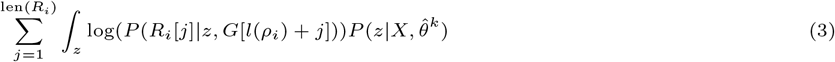

## Discussion

We have improved the ARYANA aligner as the ARYANA-BS aligner to align WGBS reads. ARYANA-BS uses the theoretical knowledge about the methylation ratio of each position of the genome to have a better alignment.

We have compared ARYANA-BS with BSMAP, bwa-meth, Bismark, BSBolt, and abismal to assess the proposed algorithm. The comparison has shown that the ARYANA-BS aligner improves previous works in the amount of correctly mapped reads for simulated reads and reduces penalties and increases mapped reads in the alignment of real reads. Additionally, ARYANA-BS has a comparable memory footprint and runtime to other tools.

ARYANA-BS is more resistant to biases in the genomic context, such as hypermethylation, hypomethylation, indels and SNP counts. This makes it the best choice for analyses that are based on genomic changes, such as cancer analysis and cell-free DNA analysis.

## Acknowledgments

The authors are grateful to Maryam Rabiee who made great contributions to the initial development of ARYANA-BS. We thank Richard Dannebaum for his support in this work.

## Notes

### Competing Interest Statement

The authors have declared no competing interest.

